# Modelling how the altered usage of cell entry pathways by the SARS-CoV-2 Omicron variant may affect the efficacy and synergy of TMPRSS2 and Cathepsin B/L inhibitors

**DOI:** 10.1101/2022.01.13.476267

**Authors:** Pranesh Padmanabhan, Narendra M. Dixit

## Abstract

The SARS-CoV-2 Omicron variant harbours mutations in its spike protein, which may affect its cell entry, tropism, and response to interventions. To elucidate these effects, we developed a mathematical model of SARS-CoV-2 entry into cells and applied it to analyse recent *in vitro* data. SARS-CoV-2 enters cells using host proteases, either Cathepsin B/L or TMPRSS2. We estimated >4-fold increase and >3-fold decrease in entry efficiency using Cathepsin B/L and TMPRSS2, respectively, of the Omicron variant relative to the original or other strains in a cell type-dependent manner. Our model predicted that Cathepsin B/L inhibitors would be more and TMPRSS2 inhibitors less efficacious against the Omicron than the original strain. Furthermore, the two inhibitor classes would exhibit synergy, although the drug concentrations maximizing synergy would have to be tailored to the Omicron variant. These findings provide insights into the cell entry mechanisms of the Omicron variant and have implications for interventions.

## INTRODUCTION

The SARS-CoV-2 variant Omicron, declared a variant of concern on 25^th^ November 2021, has become the dominant strain worldwide. It harbours as many as 37 mutations in the spike protein compared to the original SARS-CoV-2 strain (Cameroni et al., 2021) and is able to evade neutralising antibodies generated by previously infected or vaccinated individuals (Cameroni et al., 2021; Garcia-Beltran et al., 2021; Hoffmann and et al, 2021; Willett et al., 2022), possibly leading to its increased transmissibility and rapid global spread. The spike protein facilitates the entry of the virus into cells (Hoffmann et al., 2020a; Koch et al., 2021). Indeed, emerging data indicates that the high number of mutations in the Omicron variant affects its viral entry properties and cell tropism (Garcia-Beltran et al., 2021; Hoffmann and et al, 2021; Meng et al., 2021; Peacock et al., 2022; Willett et al., 2022; Zhao et al., 2021), which in turn may influence its ability to establish infection post-exposure and the severity of the subsequent symptoms. How the mutations influence entry and the efficacies of entry inhibitors remains to be elucidated.

Early studies showed that the original SARS-CoV-2 (Wuhan-Hu-1) strain displayed broad cell tropism, with viral entry a key determinant of the tropism both *in vitro* (Hoffmann et al., 2020a) and *in vivo* (Liu et al., 2021). Intriguingly, multiple recent *in vitro* studies suggest that cell tropism of the Omicron variant may be altered and its entry efficiency may be different from the original and other variants in a cell line-dependent manner (Garcia-Beltran et al., 2021; Hoffmann and et al, 2021; Meng et al., 2021; Peacock et al., 2022; Zhao et al., 2021). For instance, SARS-CoV-2 pseudotyped virus bearing either the B.1 or the Delta variant spike protein showed higher entry efficiency than the Omicron pseudotyped virus in Caco-2 (human, colon) and Calu-3 (human, lung) cells, whereas the Omicron pseudotyped virus entered Vero (African green monkey, kidney) and 293T (human, kidney) cells more efficiently than the other variants(Hoffmann and et al, 2021). Similar trends were observed in live SARS-CoV-2 virus infection assays (Meng et al., 2021; Zhao et al., 2021): the Delta variant infection spread was significantly greater than the Omicron variant in Calu-3 cells (Meng et al., 2021; Zhao et al., 2021), whereas the spread of the two variants was similar in VeroE6 cells (Zhao et al., 2021). What causes the entry efficiency of the Omicron variant relative to other variants to be higher in some cells and lower in others?

The first step in SARS-CoV-2 entry into target cells is the binding of the viral spike protein, S, with the host cell surface receptor angiotensin-converting enzyme 2 (ACE2) (Balistreri et al., 2021; Jackson et al., 2022). Cell tropism is thus expected to be affected by ACE2 expression levels (Hoffmann et al., 2020a; Koch et al., 2021; Liu et al., 2021). The Omicron spike protein binds soluble human ACE2 strongly (Cameroni et al., 2021; Hoffmann and et al, 2021). Further, ACE2 is necessary for the entry of all SARS-CoV-2 variants (Garcia-Beltran et al., 2021; Hoffmann and et al, 2021; Hoffmann et al., 2020a; Meng et al., 2021; Zhao et al., 2021). Thus, any variation in ACE2 expression across cell types is likely to have similar effects on the entry efficiencies of all the variants. The cell type-dependent variation in the entry efficiency of the Omicron variant is therefore unlikely to arise from the variations in the ACE2 expression level across cell types. Omicron spike protein incorporation into pseudotyped virus appears to be compromised compared to the Delta and Wuhan D614G strains (Meng et al., 2021). However, this reduction in spike protein density is expected to decrease the Omicron entry efficiency across all cell types, thus ruling it out as a potential cause of the differential entry efficiency observed.

Following ACE2 engagement, successful entry requires that the spike protein be cleaved into subunits by host proteases, either by the transmembrane serine protease TMPRSS2 at the plasma membrane or the cysteine proteases Cathepsin B and Cathepsin L in the endosomal vesicles (Jackson et al., 2022). The proteases TMPRSS2 and Cathepsin B/L are thought to work independently, facilitating SARS-CoV-2 entry through two independent pathways (Koch et al., 2021; Padmanabhan et al., 2020) (Fig. 1A). We recently analysed data on SARS-CoV-2 pseudotyped virus infection *in vitro* (Hoffmann et al., 2020a) using a mathematical model of SARS-CoV-2 entry and found that the relative usage of the TMPRSS2 and Cathepsin B/L entry pathways by the original SARS-CoV-2 strain varied widely across cell lines (Padmanabhan et al., 2020). For example, Vero cells predominantly admitted entry through the Cathepsin B/L pathway, whereas Calu-3 cells allowed entry via the TMPRSS2 pathway. Vero cells overexpressing TMPRSS2 permitted entry via both pathways (Hoffmann et al., 2020a). Importantly, the original strain displayed the ability to use either pathway, with the preferred pathway possibly based on the relative expression levels of the two proteases (Koch et al., 2021). We reasoned that the Omicron variant might use these pathways differently from the original (or the Delta) strain, possibly underlying its differential entry efficiency and altered cell tropism.

**Figure 1.**
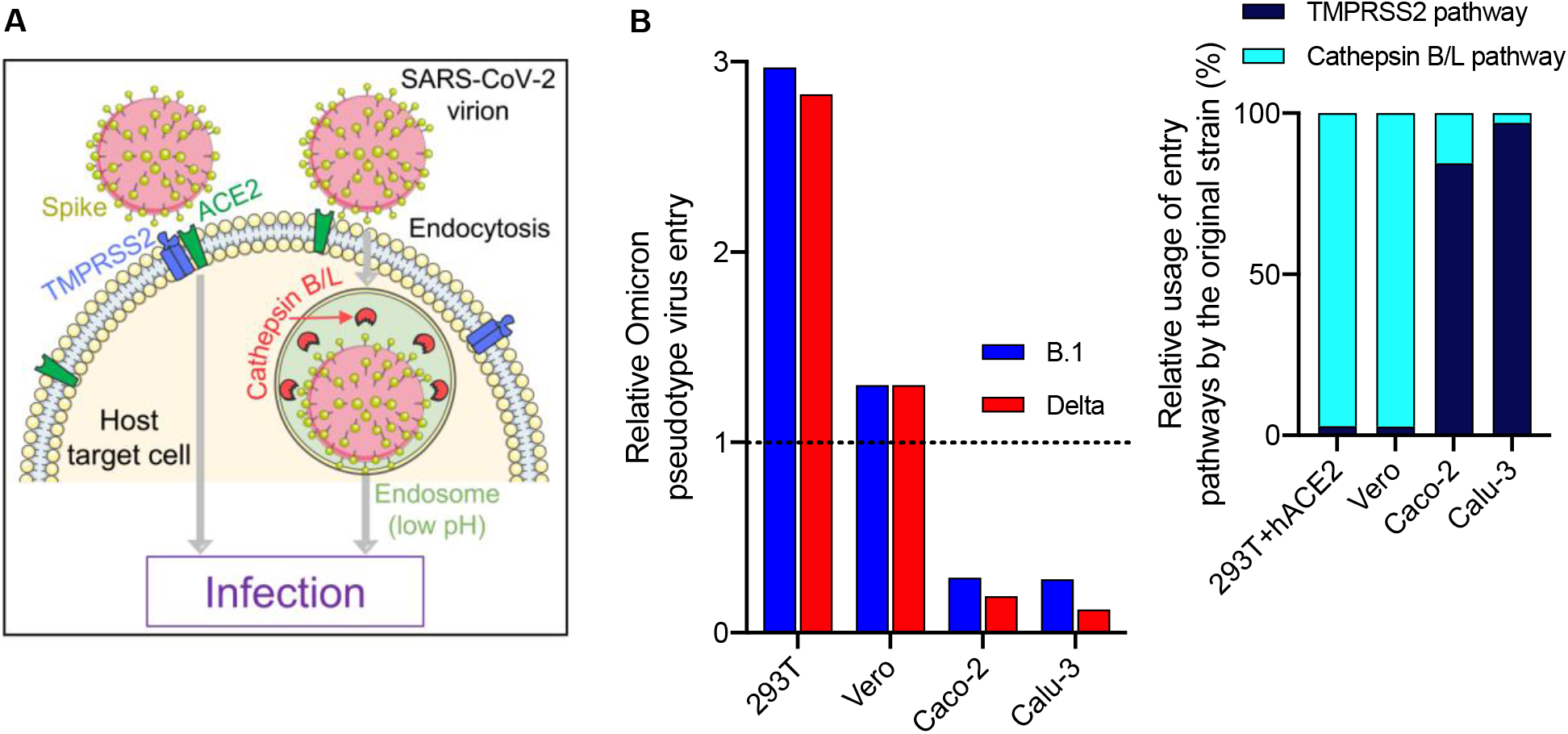
Omicron variant entry efficiency relative to B.1 and Delta strains correlates with the relative usage of the TMPRSS2 and Cathepsin B/L pathways. **(A)** Schematic of the two independent entry pathways accessible to SARS-CoV-2. **(B)** The Omicron entry efficiency relative to either the B.1 (blue) or the Delta (red) variant across cell types. Data was taken from Hoffmann et al. (2021). *Inset*, the relative usage of the entry pathways by the original strain. Data was taken from Padmanabhan et al. (2020).

Here, to elucidate the entry mechanisms of the Omicron variant, we developed a mathematical model of SARS-CoV-2 entry that explicitly considered the two independent entry pathways. Applying the model to the analysis of available *in vitro* data, we deduced the usage of the two pathways by the Omicron variant relative to the other strains. We then applied the model to predict the efficacies of drugs that target host proteases facilitating entry via the two pathways.

## METHODS

### Model of SARS-CoV-2 entry and the efficacy of entry inhibitors targeting host proteases

*In vitro* experiments typically employ virions or pseudotyped viral particles bearing the spike protein, S, of a specific SARS-CoV-2 strain to assess its entry efficiency, determined by measuring the fraction of cells infected post virus exposure. We previously developed a mathematical model to analyse such experiments (Padmanabhan et al., 2020). We adapted the model here to compare the pathway usage of the Omicron variant relative to other variants and to predict the effect of entry inhibitors.

In culture, the expression levels of host proteases are expected to vary across cells and affect viral entry into the cells. We let the expression levels, *n*_*t*_ and *n*_*c*_, of TMPRSS2 and Cathepsin B/L, respectively, follow the log-normal distributions 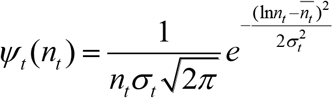 and 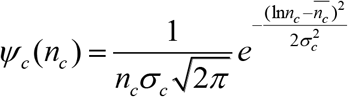, with 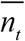 and 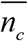 the associated means, and *σ*_*t*_ and *σ*_*c*_ the standard deviations. Thus, at the start of infection, a fraction *ϕ*_*tc*_ =*ψ*_*t*_ (*n*_*t*_)*ψ*_*c*_ (*n*_*c*_)Δ*n*_*t*_ Δ*n*_*c*_, of the cells in culture would express proteases in the narrow range Δ*n*_*t*_ and Δ*n*_*c*_ around *n*_*t*_ and *n*_*c*_, respectively. Recognizing that entry efficiency would increase with protease expression (Padmanabhan et al., 2020; Padmanabhan and Dixit, 2011, 2012), we defined the relative susceptibility of these latter cells to virus entry through the TMPRSS2 pathway, *S*_*t*_, and the Cathepsin B/L pathway, *S*_*c*_, as 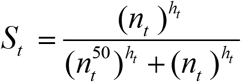 and 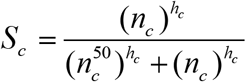, where *h*_*t*_ and *h*_*c*_ were Hill coefficients, and 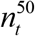 and 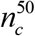 were expression levels at which *S*_*t*_ = 0.5 and *S*_*c*_ = 0.5, respectively. The overall susceptibility of the cells was thus *S*_*tc*_ = *S*_*t*_ *+ S*_*c*_ − *S*_*t*_ *S*_*c*_, based on the independence of the two pathways.

We defined *T*_*tc*_ as the above subpopulation (or fraction) of cells and considered a total of *M* × *N* such subpopulations, with *t* = 1, 2,.., *M* and *c* = 1, 2,.., *N*, defining the range of expression levels of the two proteases across cells. The following equations then described the ensuing viral dynamics:

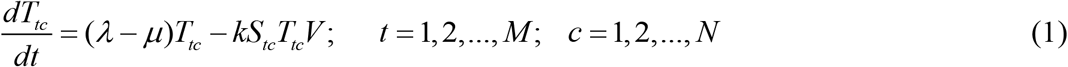

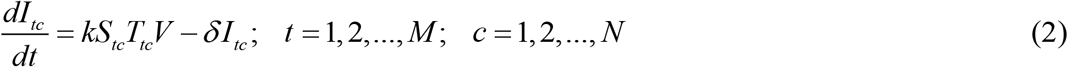

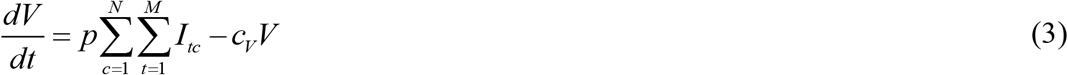

Here, cells in the subpopulation *T*_*tc*_ proliferate and die with rate constants *λ* and *µ*, respectively, and get infected by free virions, *V*, with the second order rate constant *kS*_*tc*_. *k* is thus the rate constant of the infection of the subpopulation for which entry is not limited by the proteases, *i*.*e*., for cells with *S*_*tc*_ =1. Infection produces corresponding infected cell subpopulations *I*_*tc*_, which release free virions at the rate *p* per cell and die with a rate constant *δ*. Virions are cleared with the rate constant *c*_*V*_. With pseudotyped viruses, which are replication incompetent, we let *p* = 0.

In the presence of a TMPRSS2 inhibitor, a Cathepsin B/L inhibitor, or both, the susceptibilities were lowered to *S*_*tc*_ (*D*_*T*_), *S*_*tc*_ (*D*_*C*_), and *S*_*tc*_ (*D*_*T*_, *D*_*C*_), respectively, where *D*_*T*_ and *D*_*C*_ were the concentrations of the TMPRSS2 and Cathepsin B/L inhibitors. We describe the latter susceptibilities below.

### Effect of host protease inhibitors

In the presence of a protease inhibitor targeting a specific protease, the drug would bind and block a fraction of the protease molecules from facilitating entry. With a TMPRSS2 inhibitor at concentration *D*_*T*_, we described the abundance of free TMPRSS2, denoted 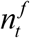, using 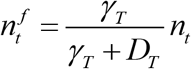, where *γ*_*T*_ was the drug concentration that reduced the abundance by half. The expression for 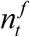 could be derived mechanistically assuming reaction equilibrium of drug-protease binding and species balance constraints (Padmanabhan et al., 2020; Padmanabhan and Dixit, 2011). The susceptibility of cells *T*_*tc*_ to infection through the TMPRSS2 pathway thus reduced to 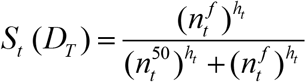. We assumed that the susceptibility through the Cathepsin B/L pathway was unaffected by a TMPRSS2 inhibitor. Similarly, in the presence of a Cathepsin B/L inhibitor at concentration *D*_*C*_, we let the abundance of free Cathepsin B/L, 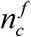, follow 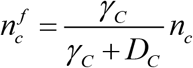, and the susceptibility through the Cathepsin B/L pathway be reduced to 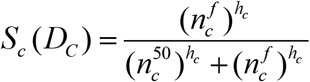. When both types of inhibitors are used simultaneously, the susceptibility was *S*_*tc*_ (*D*_*T*_, *D*_*C*_) = *S*_*t*_ (*D*_*T*_) + *S*_*c*_ (*D*_*C*_) − *S*_*t*_ (*D*_*T*_)*S*_*c*_ (*D*_*C*_).

To assess whether the drugs exhibited synergy, we predicted the population of infected cells in the absence of drugs, *I*_*tc*_, in the presence of a TMPRSS2 inhibitor, *I*_*tc*_ (*D*_*T*_), a Cathepsin B/L inhibitor, *I*_*tc*_ (*D*_*C*_), and both, *I*_*tc*_ (*D*_*T*_, *D*_*C*_), at any given time following the start of infection. The total fractions of cells unaffected by the drugs individually and together were then given by 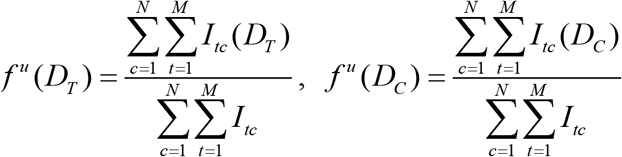, and 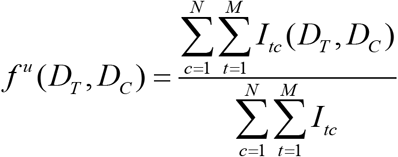.

We computed the expected fraction of cells unaffected by drugs assuming Bliss independence using 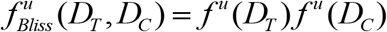. The extent of Bliss synergy then followed as 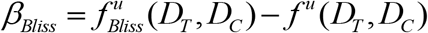, so that

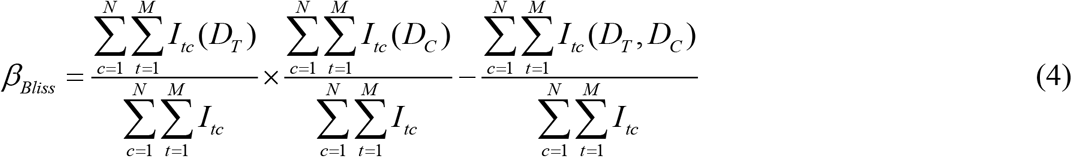

### Mean-field model to compare entry efficiencies of variants

To quantify the relative efficiency of the usage of the Cathepsin B/L and TMPRSS2 pathways by the Omicron variant, we developed a mean-field version of the model above by considering ‘average’ susceptibilities of the cells to entry via the two pathways. We thus let target cells, *T*, be infected by pseudotyped virions, *V*, to produce infected cells, *I*, depending on the ‘mean’ expression levels of TMPRSS2 and Cathepsin B/L. The following equations then described the ensuing dynamics:

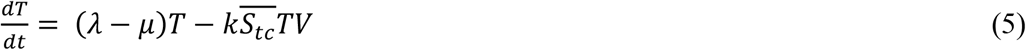

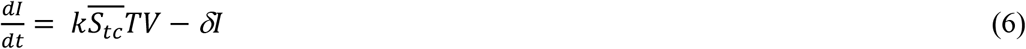

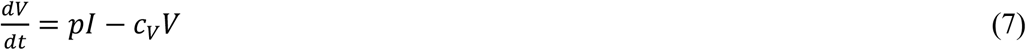

Here, we defined the average susceptibility of the cell population to entry via the Cathepsin B/L pathway as 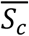 and that via the TMPRSS2 pathway as 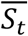. These susceptibilities increase with the expression levels of the respective proteases and saturate to unity when the levels are in excess. For a given cell type, a viral variant that has a higher 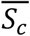 (or 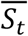) would be more efficient at entry via the Cathepsin B/L (or TMPRSS2) pathway than a variant with a lower 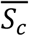 (or 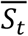). Assuming entry via the two pathways to be independent, the overall susceptibility of the population to entry can be written as 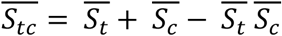. This susceptibility determined the rate of target cell infection relative to the maximal rate *k*. We let *p* = 0 as described above.

### Data analysis using the mean-field model

Experiments with pseudotyped virions typically last less than 24 h. It is reasonable to assume that cell proliferation and death do not affect the infection dynamics significantly during this period. Therefore, we ignored the proliferation and death terms in Eqs. (5-6), which allowed us to derive an analytical expression for the time evolution of the population of infected cells: 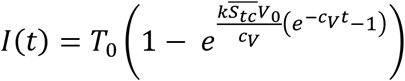, where *T*_0_ and *V*_0_ are the initial target cell density and viral concentration, respectively. Using this expression, we obtained the fraction of cells infected by the pseudotyped virions, *χ*, at time *t*_*d*_ post-infection as

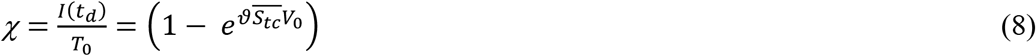

where 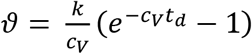 is a constant for a given *t*_*d*_.

We fit this expression to measured data from recent assays and estimated the efficiency of the usage of the two pathways by the Omicron variant relative to the original or other strains.

### Model parameters

We chose model parameters, such as the target cell proliferation and death rate constants, from the literature (Gonçalves et al., 2020; Hoffmann et al., 2020a; Jiang et al., 2019; Padmanabhan and Dixit, 2015; Ursache et al., 2015) and our previous study (Padmanabhan et al., 2020). The model parameter values and initial conditions are summarised in Table 1.

**Table 1:**
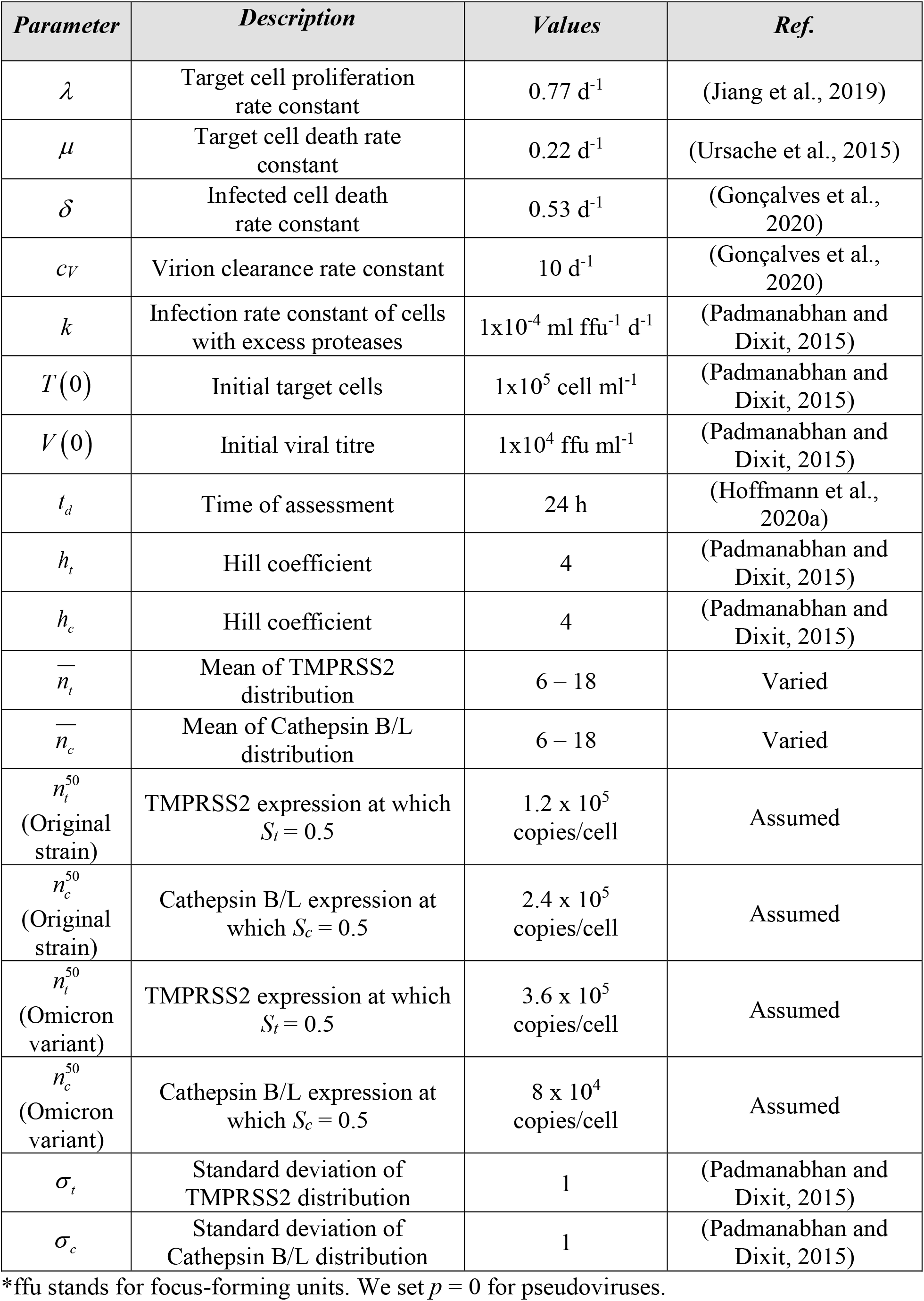
Model parameters and their values.

| *Parameter* | *Description* | *Values* | *Ref.* |
| --- | --- | --- | --- |
| $\lambda$ | Target cell proliferation rate constant | $0.77 \text{ d}^{-1}$ | (Jiang et al., 2019) |
| $\mu$ | Target cell death rate constant | $0.22 \text{ d}^{-1}$ | (Ursache et al., 2015) |
| $\delta$ | Infected cell death rate constant | $0.53 \text{ d}^{-1}$ | (Gonçalves et al., 2020) |
| $c_V$ | Virion clearance rate constant | $10 \text{ d}^{-1}$ | (Gonçalves et al., 2020) |
| $k$ | Infection rate constant of cells with excess proteases | $1 \times 10^{-4} \text{ ml ffu}^{-1} \text{ d}^{-1}$ | (Padmanabhan and Dixit, 2015) |
| $T(0)$ | Initial target cells | $1 \times 10^5 \text{ cell ml}^{-1}$ | (Padmanabhan and Dixit, 2015) |
| $V(0)$ | Initial viral titre | $1 \times 10^4 \text{ ffu ml}^{-1}$ | (Padmanabhan and Dixit, 2015) |
| $t_d$ | Time of assessment | 24 h | (Hoffmann et al., 2020a) |
| $h_t$ | Hill coefficient | 4 | (Padmanabhan and Dixit, 2015) |
| $h_c$ | Hill coefficient | 4 | (Padmanabhan and Dixit, 2015) |
| $\bar{n}_t$ | Mean of TMPRSS2 distribution | 6 – 18 | Varied |
| $\bar{n}_c$ | Mean of Cathepsin B/L distribution | 6 – 18 | Varied |
| $n_t^{50}$<br>(Original strain) | TMPRSS2 expression at which $S_t = 0.5$ | $1.2 \times 10^5$ copies/cell | Assumed |
| $n_c^{50}$<br>(Original strain) | Cathepsin B/L expression at which $S_c = 0.5$ | $2.4 \times 10^5$ copies/cell | Assumed |
| $n_t^{50}$<br>(Omicron variant) | TMPRSS2 expression at which $S_t = 0.5$ | $3.6 \times 10^5$ copies/cell | Assumed |
| $n_c^{50}$<br>(Omicron variant) | Cathepsin B/L expression at which $S_c = 0.5$ | $8 \times 10^4$ copies/cell | Assumed |
| $\sigma_t$ | Standard deviation of TMPRSS2 distribution | 1 | (Padmanabhan and Dixit, 2015) |
| $\sigma_c$ | Standard deviation of Cathepsin B/L distribution | 1 | (Padmanabhan and Dixit, 2015) |

645 \*ffu stands for focus-forming units. We set $p = 0$ for pseudoviruses.

## RESULTS

### Correlation of entry efficiency with pathway usage

To test our hypothesis that the Omicron variant uses the entry pathways differently, we examined data from the recent experiments of Hoffmann et al. (2021), who measured the extent of infection of cells *in vitro* by the Omicron variant (BA.1 sublineage) relative to other variants in different cell types. Interestingly, we found a correlation between the entry efficiency of the Omicron variant and the relative usage of the two entry pathways by the original strain (Fig. 1B). With cell lines (Calu-3 and Caco-2) where the usage of the TMPRSS2 entry pathway by the original strain was dominant, the Omicron pseudotyped virus entry was significantly less efficient than the B.1 and delta strains. By contrast, the Omicron virus entry was more efficient in cell lines (293T and Vero) where the Cathepsin B/L entry pathway usage by the original strain was dominant (Fig. 1B). One possible interpretation of this relationship is that the Omicron variant entry is relatively less efficient through the TMPRSS2 pathway and more efficient via the Cathepsin B/L pathway than the other strains. Consistent with this notion, camostat mesylate, a TMPRSS2 inhibitor, was less potent against the Omicron variant than the Delta variant in blocking live virus infection of VeroE6 cells overexpressing TMPRSS2, which allows entry via both pathways (Zhao et al., 2021). Moreover, syncytium formation in a cell-cell fusion assay, which requires TMPRSS2 but not Cathepsins, was severely impaired for the Omicron variant compared to both the Delta (Meng et al., 2021; Zhao et al., 2021) and the Wuhan-Hu-1 D614G (Meng et al., 2021) strains. The increased efficiency of Cathepsin B/L usage together with the reduced efficiency of TMPRSS2 usage by the Omicron variant may explain its altered cell tropism relative to the original strain or other variants.

### Omicron variant entry with low Cathepsin B/L expression

We reasoned that the increased efficiency of Cathepsin B/L usage and reduced TMPRSS2 usage by the Omicron variant would imply that the variant would successfully enter cells that express low Cathepsin B/L levels, where the original strain might fail. To elucidate this, we performed calculations using our model (Eqs. (1-3)) by varying the expression level of Cathepsin B/L. We first set 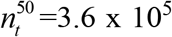 copies/cell and 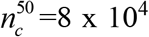 copies/cell for the Omicron variant and 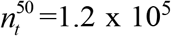 copies/cell and 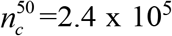 copies/cell for the original strain so that for given distributions of the TMPRSS2 and Cathepsin B/L expression levels, the susceptibility of cells to entry via the TMPRSS2 pathway was lower (Fig. 2A) and the Cathepsin B/L pathway was higher (Fig. 2B) for the Omicron variant than the original strain. Recall that 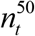 is the expression level at which entry efficiency via the TMPRSS2 pathway is half-maximal. Thus, the lower the 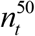, the more efficient is the usage of the TMPRSS2 pathway. Analogous notions apply to the Cathepsin B/L pathway. We then considered three *in silico* cell lines with a constant TMPRSS2 distribution (Fig. 2C) but different Cathepsin B/L levels, the latter termed low, medium and high (Fig. 2D). For each of these cell lines, we predicted the distributions of cells infected by the original strain as well as by the Omicron variant (Fig. 2E) and compared the two (Fig. 2F). We found that when Cathepsin B/L expression was high, the advantage of the Omicron variant was undermined because of the abundance of Cathepsin B/L, so that the two strains had similar entry levels (Fig. 2F). As the expression level decreased, the Omicron variant was more successful at entry. With medium expression level, the Omicron variant had ∼1.4-fold greater entry, and with low expression level, it had more than ∼2.9-fold greater entry success than the original strain.

**Figure 2.**
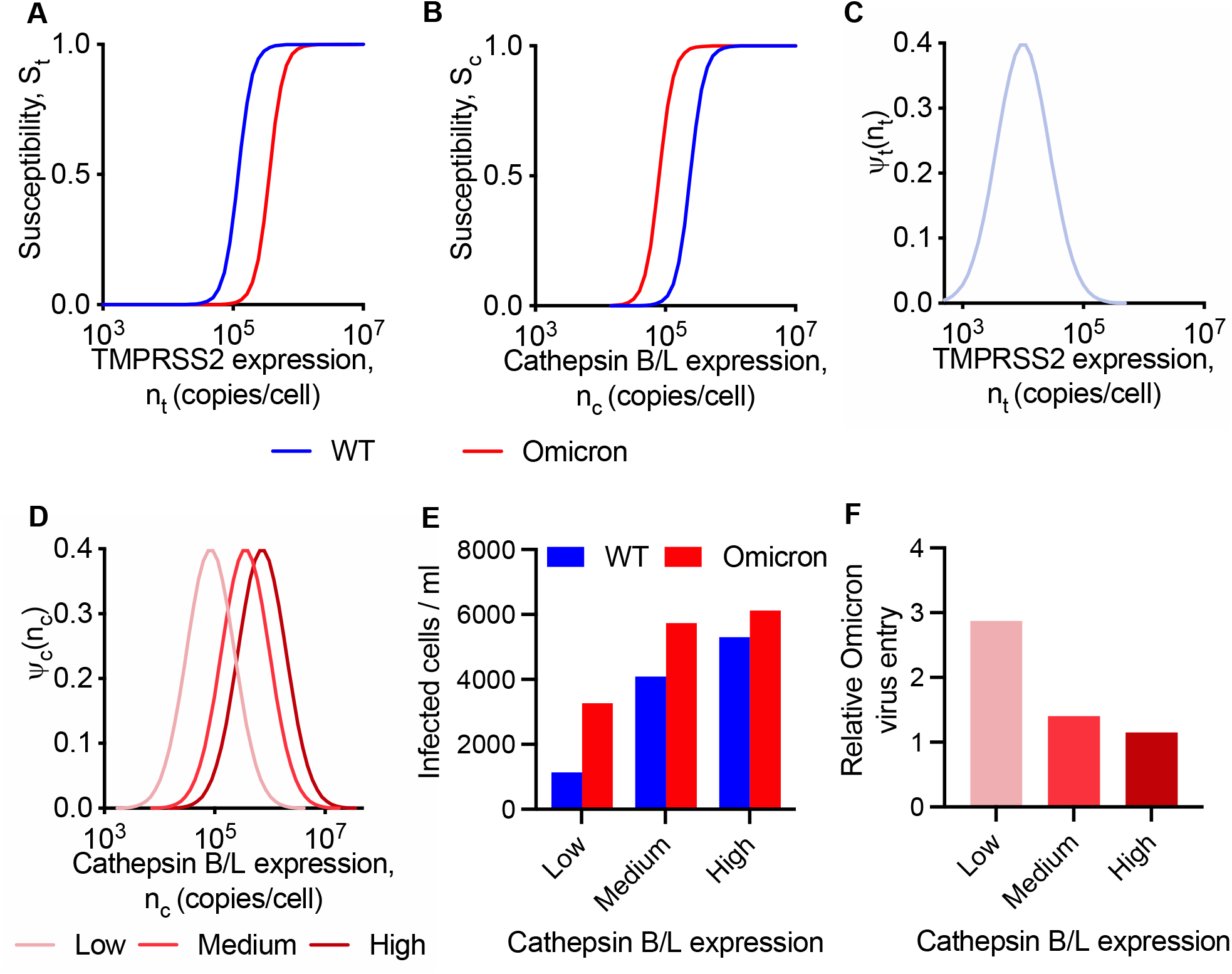
The relative entry efficiency of the Omicron variant depends on the protease expression. **(A, B)** Dependence of susceptibility of infection on TMPRSS2 (A) and Cathepsin B/L (B) expression levels for the original strain (blue line) and Omicron variant (red line). **(C)** The log-normal distribution of TMPRSS2 across cells with the mean TMPRSS2 expression levels 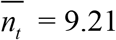. **(D)** The log-normal distribution of Cathepsin B/L across cells with low 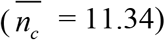, medium 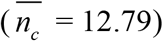 and high 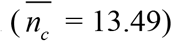 mean Cathepsin B/L expression levels. **(E, F)** Cells infected by the wild-type (blue bar) and Omicron variant (red bar) (E) and the Omicron entry efficiency relative to the original strain (F) at different Cathepsin B/L expression levels shown in Fig. 2D and fixed TMPRSS2 expression level shown in Fig. 2C. The other parameters and initial conditions are listed in Table 1.

With these predictions further strengthening our hypothesis above, we asked whether we could apply our model to analyse available data to quantify the increased (decreased) entry efficiency of the Omicron variant via the Cathepsin B/L (TMPRSS2) pathway. Because the distributions of the protease levels across cells are not known, we employed a simplified, mean-field version of our model for this analysis (Methods).

### Quantitative estimates of relative entry efficiency

We considered data from cell types where the original strain nearly exclusively entered via either the Cathepsin B/L or the TMPRSS2 pathways, so that we could estimate the relative efficiency of the usage of either pathway by the Omicron variant. We thus examined first data from Garcia-Beltran et al. (2021), who measured the fraction of 293T-ACE2 cells infected by pseudotyped virus bearing the wild-type or the Omicron spike proteins at different viral concentrations, *V*_0_ (Fig. 3A). 293T-ACE2 cells predominantly permit original strain entry via the Cathepsin B/L pathway (Padmanabhan et al., 2020). Given the relatively poor efficiency of TMPRSS2 usage by the Omicron variant, we made the approximation 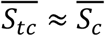 in Eqs. (5-6). Using the resulting expression (Eq. (8)), we fit predictions of *χ* as a function of *V*_0_ to the corresponding experimental data (Garcia-Beltran et al., 2021) and estimated the composite parameter 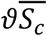 for the wild-type (wt) and the Omicron variant. Our model provided good fits to the data (Fig. 3A). Taking ratios of the estimated composite parameter and recognizing that *ϑ* is unlikely to be strain dependent, we obtained 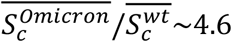, indicating an ∼4.6-fold increased efficiency of the usage of the Cathepsin B/L pathway by the Omicron variant in 293T-ACE2 cells relative to the original strain.

**Figure 3.**
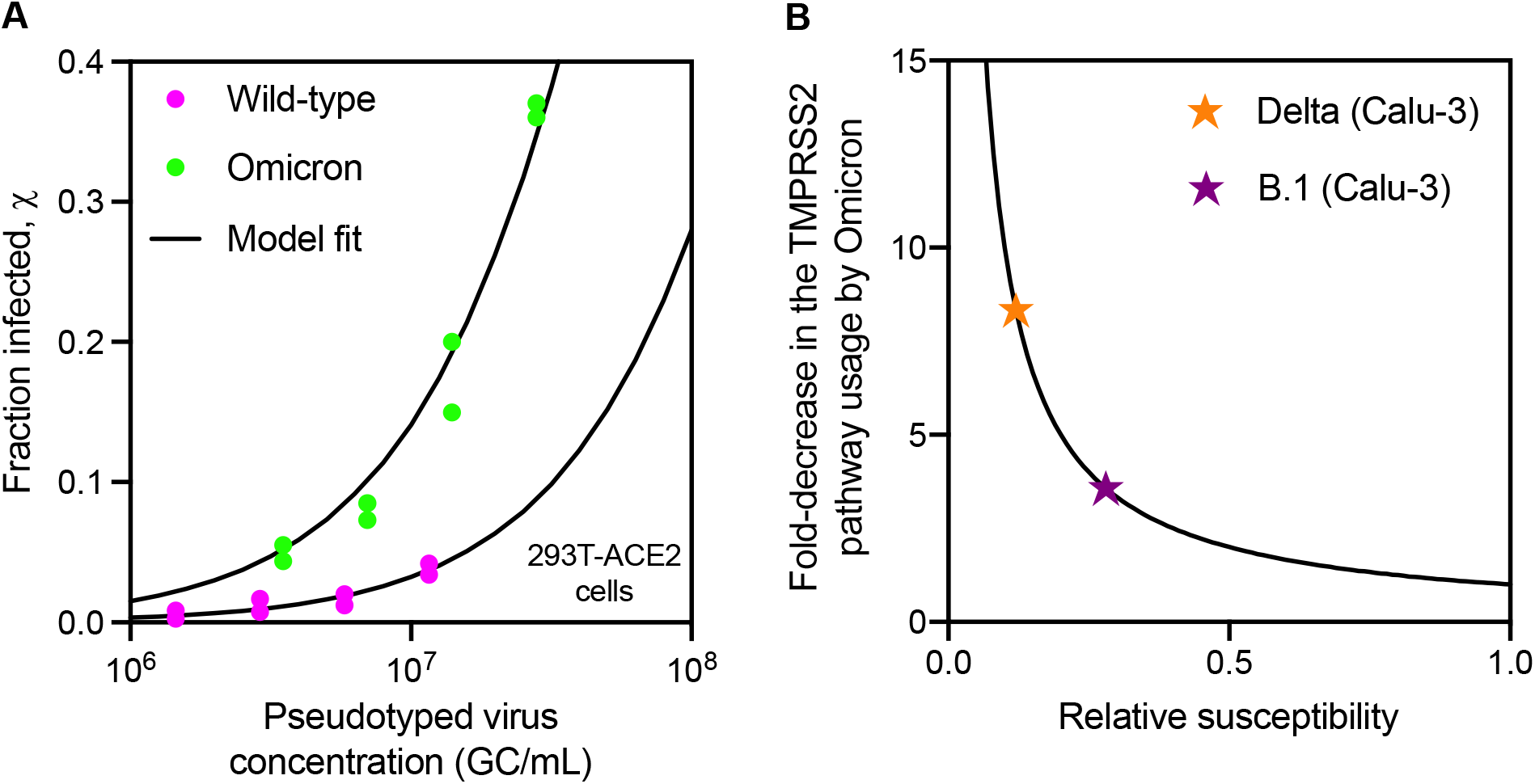
Estimation of entry pathway usage by the Omicron variant. **(A)** Fits of model predictions (lines) to the experimental data (symbols) taken from Garcia-Beltran et al. (2021) measuring the fraction of cells infected by pseudotyped virus bearing either the wild-type or the Omicron spike proteins. Data was fit using the tool NLINFIT in MATLAB R2017b. **(B)** The fold-decrease in Omicron entry efficiency through the TMPRSS2 pathway as a function of relative susceptibility to entry. Symbols place experimental data of relative infection taken from Hoffmann et al. (2021) on the relative susceptibility curve predicted (see text) so that the fold decrease can be read off. In A and B, experimental data was extracted using Engauge Digitizer 12.1.

We next analysed the experiments of Hoffmann et al. (2021) mentioned above (Fig 1B). We focussed on Calu-3 cells, which appear to allow entry of the original strain nearly exclusively via TMPRSS2. Here, we therefore made the approximation 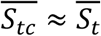 in Eqs. (5-6). Unlike the above dataset, measurements here were available at a single initial viral load. Further, measurements were available of the fractions of cells infected by viruses pseudotyped with the Omicron variant relative to those with the B.1 or Delta variants. Note that B.1 has spike proteins identical to the original strain except for the D614G mutation. Because the overall infection of Calu-3 cells is small, Taylor series expansion of Eq. (7) yielded the approximation 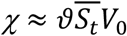. Accordingly, the susceptibility of cells to entry via the TMPRSS2 pathway of the Omicron variant relative to the B.1 strain would be given by the ratio of the fractions of cells infected in the two assays: 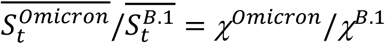. Using the latter expression and the corresponding data yielded 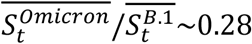 and 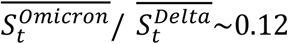. Thus, we estimated ∼3.6-fold and ∼8.3-fold decreased efficiency of the Omicron variant in using TMPRSS2 for entry in Calu-3 cells compared to the B.1 and Delta strains, respectively (Fig. 3B).

Together, thus, the Omicron variant appears to have evolved to use Cathepsin B/L more efficiently and TMPRSS2 less efficiently for virus entry than the original and other strains.

### Efficacies of Cathepsin B/L and TMPRSS2 inhibitors in blocking Omicron variant entry

Drugs targeting the proteases Cathepsin B/L and TMPRSS2 offer promising routes to treat SARS-CoV-2 infection (Hashimoto et al., 2021; Hoffmann et al., 2020a; Hoffmann et al., 2020b; Kreutzberger et al., 2021; Ou et al., 2021). The improved efficiency of Cathepsin B/L usage and reduced efficiency of TMPRSS2 usage would imply that drugs targeting the former but not the latter would be preferred for treating infections with the Omicron variant. To examine this, we applied our model (Eqs. (1-3)) to predict the ability of the two classes of inhibitors to prevent viral entry. We considered an *in silico* cell line with log-normal distributions of the expression levels of TMPRSS2 and Cathepsin B/L (Fig. 4A) and the strain-dependent susceptibility of cells to entry via the two pathways (Fig. 2A, B). In the presence of a TMPRSS2 inhibitor, our model predicted that entry of the original strain would decrease in a dose-dependent manner and saturate, for the parameters chosen, to ∼45% of that in the absence of the drug (Fig. 4B). The latter plateau represented the entry via the Cathepsin B/L pathway when entry via the TMPRSS2 pathway was fully blocked. Under the same conditions, entry of the Omicron variant saw hardly any decrease compared to that in the absence of the drug. This is because the Omicron variant entry proceeded only minimally via the TMPRSS2 pathway, leaving little room for the drug to act.

**Figure 4.**
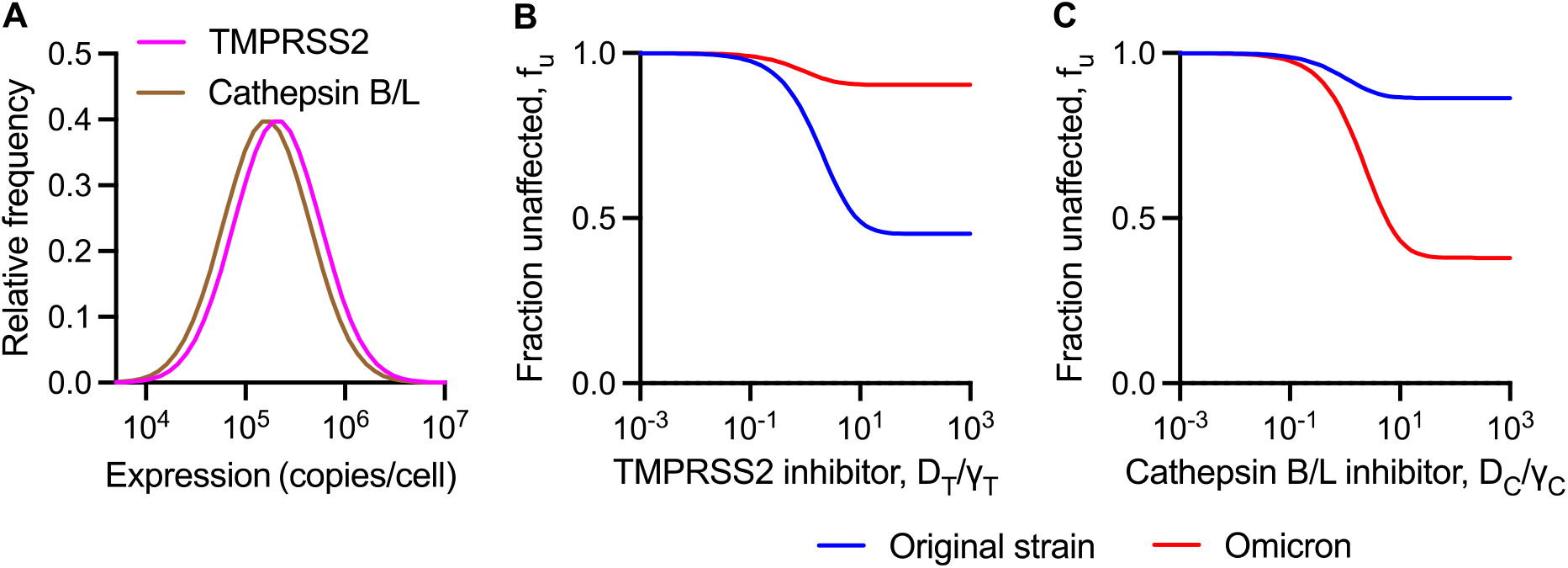
Predictions of the efficacies of TMPRSS2 and Cathepsin B/L inhibitors against the wild-type and Omicron variant. **(A)** The log-normal distribution of TMPRSS2 and cathepsin B/L across cells. **(B, C)** The fraction of infection events caused by the original strain (blue line) and Omicron variant (red line) uninhibited by different concentrations of a TMPRSS2 inhibitor (B) and a Cathepsin B/L inhibitor (C). In A-C, 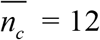 and 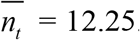. In D-E, 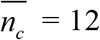. In F-G, 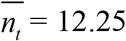. The other parameters and initial conditions are listed in Table 1.

The use of a Cathepsin B/L inhibitor, however, had the opposite effect. The entry of the original strain showed hardly any reduction with increase in drug concentration (Fig. 4C). On the other hand, the entry of the Omicron variant decreased in a dose-dependent manner and plateaued to a value of ∼38% of that in the absence of the drug.

It followed, thus, that Cathepsin B/L inhibitors would be more effective in blocking Omicron variant entry than TMPRSS2 inhibitors.

### Synergy between Cathepsin B/L and TMPRSS2 inhibitors against both the original strain and Omicron variant

Previously, we predicted synergy between the two types of inhibitors in blocking virus entry, which has since been observed in experiments with the original strain (Hoffmann et al., 2020a; Kreutzberger et al., 2021). Here, we applied our model to estimate the extent of synergy with the Omicron variant relative to the original strain (Eqs. (1-4)) (Fig. 5A-J). Given the substantial cell-type dependence in entry efficiency, we first considered two *in silico* cell types, one with relatively low (Fig. 5A) and the other with relatively high (Fig. 5E) mean TMPRSS2 expression levels. Both the cell types expressed moderate Cathepsin B/L levels (Fig. 5A, E). The TMPRSS2 and Cathepsin B/L inhibitors worked synergistically to block original strain and Omicron variant entry in a dose-dependent manner, albeit to different extents (Fig. 5B-D, F-H).

**Figure 5.**
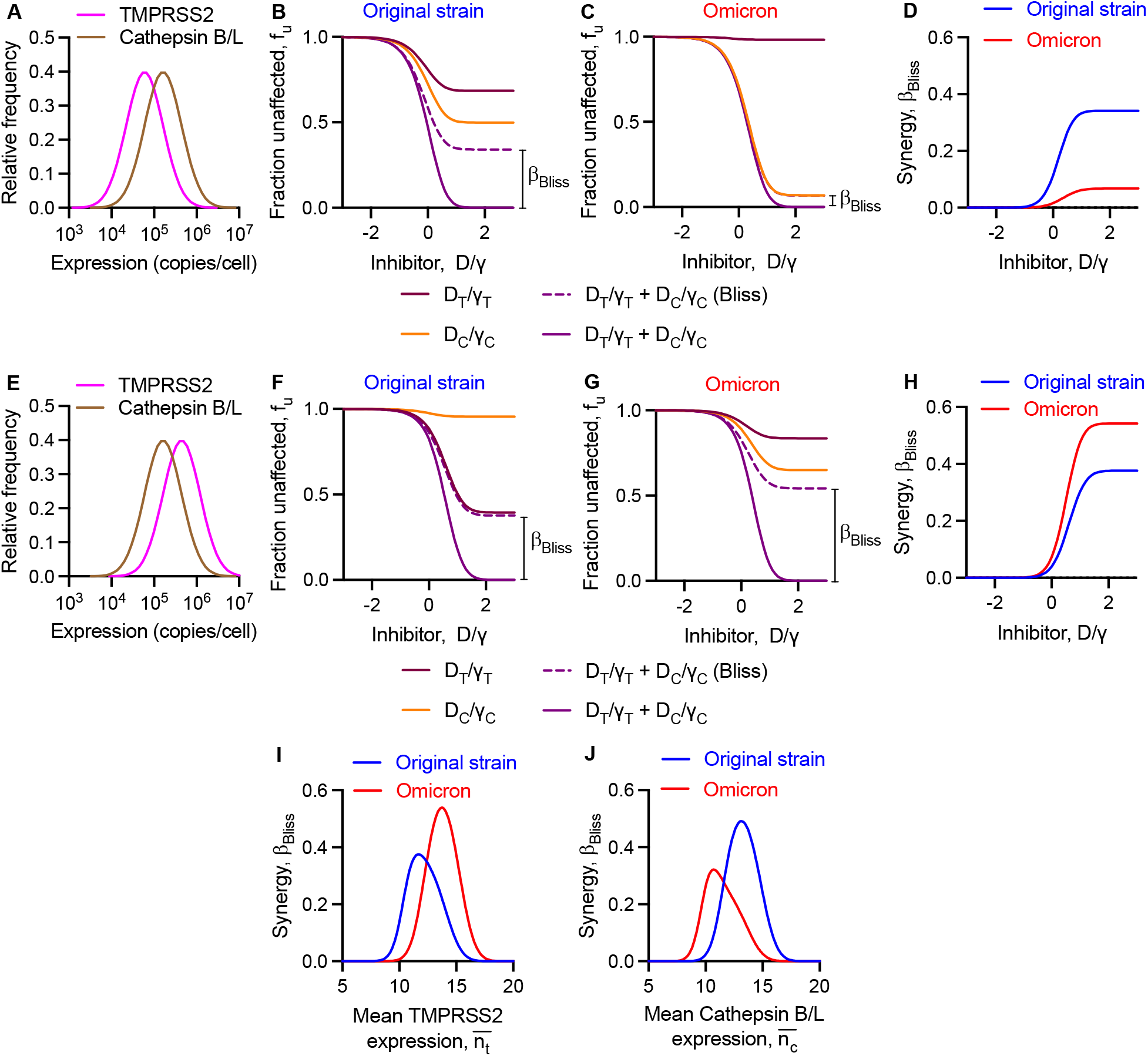
Predictions of the effect of combination treatment targeting both TMPRSS2 and Cathepsin B/L pathways against the original strain and Omicron variant. **(A-D)** The log-normal distribution of TMPRSS2 and cathepsin B/L across cells (A) employed to predict the fraction of infection events caused by the original strain (B) and the Omicron variant (C) unaffected by Cathepsin B/L inhibitor, TMPRSS2 inhibitor or both and the Bliss synergy (D) over a range of drug concentrations. **(E-H)** The log-normal distribution of TMPRSS2 and cathepsin B/L across cells (E) employed to predict the fraction of infection events caused by the original strain (F) and the Omicron variant (G) unaffected by Cathepsin B/L inhibitor, TMPRSS2 inhibitor or both and the Bliss synergy (H) over a range of drug concentrations. In B, C, F, and G, the extent of Bliss synergy is marked. **(I, J)** The predicted Bliss synergy for varying TMPRSS2 expression and fixed mean 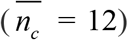 Cathepsin B/L expression and for varying Cathepsin B/L expression and fixed mean 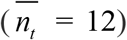 TMPRSS2 expression at two different drug concentrations. In B-D and F-H, *D*_*C*_*/γ*_*C*_ = *D*_*T*_*/γ*_*T*_ = *D*/*γ*. In I, 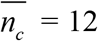. In J, 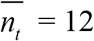. In I-J, *D*_*C*_*/γ*_*C*_ = *D*_*T*_*/γ*_*T*_ = 10. The other parameters and initial conditions are listed in Table 1.

At low drug levels, Bliss synergy, *β*_*Bliss*_, was minimal because the chances of the drugs acting together on the same cell were minimal (Fig. 5D, H). As the drug levels increased, the synergy increased and plateaued at a value that depended on the expression levels of the proteases and the viral strain (Fig. 5D, H). This effect could be understood by considering the relative usage of the two entry pathways. In our calculations, drug synergy was high when the relative preferences for the two pathways were similar, so that both pathways had to be blocked and drugs targeting both pathways would thus synergize. When usage of either of the two pathways dominated, the drug that blocked the dominant pathway alone had a role, leaving little room for synergy. For instance, when TMPRSS2 levels were low (Fig. 5A), the original strain used both pathways for entry (Fig. 5B), whereas the Omicron variant predominantly used the Cathepsin B/L pathway. Consequently, the maximum synergy achieved against the original strain (*β*_*Bliss*_ = 0.34) was higher than the Omicron variant (*β*_*Bliss*_ = 0.07) (Fig. 5B). In contrast, when TMPRSS2 levels were high (Fig. 5E), the original strain predominantly used the TMPRSS2 pathway, and the Omicron variant used both the pathways for entry. The synergy for the Omicron variant (*β*_*Bliss*_ = 0.54) was now much higher than that for the original strain (*β*_*Bliss*_ = 0.38) (Fig. 5H).

We next predicted synergy by varying either the mean expression levels of TMPRSS2 (Fig. 5I) or Cathepsin B/L (Fig. 5J). The synergy with the original strain and the Omicron variant showed a non-monotonic dependence on protease expression levels. No synergy was observed at low and high protease expression levels, when virus entry through one of the two pathways was dominant (Fig. 5I, J). Synergy peaked at intermediate protease expression levels when the usage of the two entry pathways was comparable (Fig. 5I, J). Because the Omicron variant uses the TMRPSS2 pathway less efficiently and the Cathepsin B/L pathway more efficiently than the original strain, synergy for Omicron peaked at a relatively higher TMPRSS2 (Fig. 5I) and lower Cathepsin B/L (Fig. 5J) expression levels than for the original strain. Drug levels that would yield optimal synergy would thus be different for the original strain and the Omicron variant.

In summary, our model predicts that drug combinations targeting TMPRSS2 and Cathepsin B/L inhibitors would synergistically block original strain and Omicron variant entry, albeit to different extents in a protease expression level-dependent manner.

## DISCUSSION

Recent studies have reported changes in cellular tropism and cell entry properties of the Omicron variant compared to the original SARS-CoV-2 strain and other variant (Garcia-Beltran et al., 2021; Hoffmann and et al, 2021; Meng et al., 2021; Peacock et al., 2022; Willett et al., 2022; Zhao et al., 2021). Quantifying the changes in the preference of the Omicron variant for different entry routes may help evaluate the efficacies of entry inhibitors in clinical development and optimise entry inhibitor-based treatments (Gunst et al., 2021; Zhuravel et al., 2021). Mathematical models of SARS-CoV-2 kinetics have provided valuable insights into COVID-19 disease progression, drug action, and the effectiveness of vaccines and treatments (Amidei and Dobrovolny, 2022; Chatterjee et al., 2022; Desikan et al., 2021; Garg et al., 2021; Gonçalves et al., 2020; Goyal et al., 2020; Ke et al., 2021; Kissler et al., 2021; Néant et al., 2021; Padmanabhan et al., 2022; Perelson and Ke, 2020). Here, adapting a previously developed mathematical model of SARS-CoV-2 entry (Padmanabhan et al., 2020) to the analysis of *in vitro* data on variants (Garcia-Beltran et al., 2021; Hoffmann and et al, 2021), we quantified the altered usage of host proteases required for entry by the Omicron variant relative to the original strain and the Delta variant. We then applied the model to examine the influence of the latter on the efficacies and synergy of drugs targeting the host proteases mediating entry. Our analysis suggests that the Omicron variant appears to have evolved to use Cathepsin B/L more efficiently and TMPRSS2 less efficiently than the original strain and the Delta variant for cell entry. Specifically, we estimated >4-fold enhanced efficiency in the usage of the Cathepsin B/L pathway and >3-fold reduced efficiency in the usage of the TMPRSS2 pathway by the Omicron variant in the cell lines examined. The alteration in the preference of the Omicron variant for entry pathways may have clinical implications. Drugs targeting TMPRSS2, such as camostat mesylate (Hoffmann et al., 2020a), and the Cathepsin B/L pathway, such as CA-074 methyl ester (Hashimoto et al., 2021), are in development. The efficacy of TMPRSS2 inhibitors is likely to decrease and that of Cathepsin B/L inhibitors is likely to increase against the Omicron strain. Our model predictions corroborated this expectation. Available experimental data too are consistent with this expectation. Camostat mesylate, a TMPRSS2 inhibitor, worked poorly against the Omicron variant compared to the Delta variant in blocking infection *in vitro* (Zhao et al., 2021). Future studies may test whether Cathepsin B/L inhibitors would work better against the Omicron variant than the original and Delta strains. The mechanistic origins of the altered entry pathway usage by the Omicron variant remain poorly elucidated. Although molecular dynamics simulations can offer insights (Aggarwal et al., 2021; Ray et al., 2021; Zimmerman et al., 2021), the large number of mutations in the spike protein of the Omicron variant render the mechanisms difficult to unravel.

In a previous study, we predicted synergy between TMPRSS2 and Cathepsin B/L inhibitors against original SARS-CoV-2 infection (Padmanabhan et al., 2020). This synergy has since been observed experimentally (Hoffmann et al., 2020a; Kreutzberger et al., 2021). Here, we predicted that the drugs would also exhibit synergy against the Omicron variant. The extent of synergy and the drug concentrations at which the synergy would be maximum, however, are likely to be different for the Omicron variant compared to the other strains. The difference is expected because of the different entry pathway usages involved. Our formalism offers a route to identify the optimal drug concentrations using the distributions of the expression levels of the proteases and the dependent susceptibilities of cells to entry via the two pathways as inputs. While such inputs have been available for other viruses such as HIV-1 and hepatitis C virus and enabled quantitative model predictions (Brandenberg et al., 2015; Koizumi et al., 2017; Magnus et al., 2009; Mulampaka and Dixit, 2011; Padmanabhan and Dixit, 2011, 2015, 2017; Padmanabhan et al., 2014; Sen et al., 2019; Venugopal et al., 2018), they are currently not available for SARS-CoV-2. Consequently, our findings of drug efficacies and synergy, although robust, remain qualitative.

Since late 2021, more than a hundred Omicron sublineages, including BA.1.1, BA.2, BA.2.3, BA.2.75, BA.2.9, BA.2.12.1, BA.3, BA.4, and BA.5, have emerged (Xia et al., 2022). These sublineages have evolved to exhibit different transmissibility, virulence, and immune evasion (Xia et al., 2022). Whether they also have different preferences for entry routes is not fully understood. Our study has focussed on the BA.1 sublineage. Our model presents a framework to analyse the emerging *in vitro* data and quantify the changes in the entry pathways used by the other Omicron sublineages.

Implications of our findings also follow for our understanding of the cell tropism of the Omicron variant. Because of its preferred and more efficient usage of the Cathepsin B/L pathway, it is likely to preferentially infect cells expressing high levels of Cathepsin B/L. It would be interesting to test whether such cells are present in greater abundance in the upper respiratory tract, facilitating greater transmission of the Omicron variant than other pre-Omicron variants. Similarly, future studies may test whether the reduced usage of TMPRSS2 results in less severe disease compared to the other variants.

## ACKNOWLEDGEMENTS

We thank Rajat Desikan for help with Fig. 1A.

## REFERENCES

Aggarwal, A., Naskar, S., Maroli, N., Gorai, B., Dixit, N.M., Maiti, P.K., 2021. Mechanistic insights into the effects of key mutations on SARS-CoV-2 RBD-ACE2 binding. Phys Chem Chem Phys 23, 26451–26458.

Amidei, A., Dobrovolny, H.M., 2022. Estimation of virus-mediated cell fusion rate of SARS-CoV-2. Virology 575, 91–100.

Balistreri, G., Yamauchi, Y., Teesalu, T., 2021. A widespread viral entry mechanism: The C-end Rule motif-neuropilin receptor interaction. Proc Natl Acad Sci U S A 118, e2112457118.

Brandenberg, O.F., Magnus, C., Regoes, R.R., Trkola, A., 2015. The HIV-1 entry process: A stoichiometric view. Trends Microbiol 23, 763–774.

Cameroni, E., Bowen, J.E., Rosen, L.E., Saliba, C., Zepeda, S.K., Kulap, K., al., e., 2021. Broadly neutralizing antibodies overcome SARS-CoV-2 Omicron antigenic shift. Nature 602, 664–670.

Chatterjee, B., Sandhu, H.S., Dixit, N.M., 2022. Modeling recapitulates the heterogeneous outcomes of SARS-CoV-2 infection and quantifies the differences in the innate immune and CD8 T-cell responses between patients experiencing mild and severe symptoms. PLoS Pathog 18, e1010630.

Desikan, R., Padmanabhan, P., Kierzek, A.M., van der Graaf, P.H., 2021. Mechanistic models of COVID-19: Insights into disease progression, vaccines, and therapeutics. Int J Antimicrob Agents 60, 106606.

Garcia-Beltran, W.F., Denis, K.J.S., Hoelzemer, A., Lam, E.C., Nitido, A.D., Sheehan, M.L., Berrios, C., al., e., 2021. mRNA-based COVID-19 vaccine boosters induce neutralizing immunity against SARS-CoV-2 Omicron variant. Cell 185, 457–466.

Garg, A.K., Mittal, S., Padmanabhan, P., Desikan, R., Dixit, N.M., 2021. Increased B cell selection stringency in germinal centers can explain improved COVID-19 vaccine efficacies with low dose prime or delayed boost. Front Immunol 12, 776933.

Gonçalves, A., Bertrand, J., Ke, R., Comets, E., de Lamballerie, X., Malvy, D., Pizzorno, A., Terrier, O., Rosa Calatrava, M., Mentré, F., Smith, P., Perelson, A.S., Guedj, J., 2020. Timing of antiviral treatment initiation is critical to reduce SARS-CoV-2 viral load. CPT: Pharmacometrics Syst Pharmacol 9, 509–514.

Goyal, A., Cardozo-Ojeda, E.F., Schiffer, J.T., 2020. Potency and timing of antiviral therapy as determinants of duration of SARS-CoV-2 shedding and intensity of inflammatory response. Sci Adv 6, eabc7112.

Gunst, J.D., Staerke, N.B., Pahus, M.H., Kristensen, L.H., Bodilsen, J., Lohse, N., Dalgaard, L.S., Brønnum, D., Fröbert, O., Hønge, B., Johansen, I.S., Monrad, I., Erikstrup, C., Rosendal, R., Vilstrup, E., Mariager, T., Bove, D.G., Offersen, R., Shakar, S., Cajander, S., Jørgensen, N.P., Sritharan, S.S., Breining, P., Jespersen, S., Mortensen, K.L., Jensen, M.L., Kolte, L., Frattari, G.S., Larsen, C.S., Storgaard, M., Nielsen, L.P., Tolstrup, M., Sædder, E.A., Østergaard, L.J., Ngo, H.T.T., Jensen, M.H., Højen, J.F., Kjolby, M., Søgaard, O.S., 2021. Efficacy of the TMPRSS2 inhibitor camostat mesilate in patients hospitalized with Covid-19-a double-blind randomized controlled trial. EClinicalMedicine 35, 100849.

Hashimoto, R., Sakamoto, A., Deguchi, S., Yi, R., Sano, E., Hotta, A., Takahashi, K., Yamanaka, S., Takayama, K., 2021. Dual inhibition of TMPRSS2 and Cathepsin B prevents SARS-CoV-2 infection in iPS cells. Mol Ther Nucleic Acids 26, 1107–1114.

Hoffmann, M., et al, 2021. The Omicron variant is highly resistant against antibody-mediated neutralization: Implications for control of the COVID-19 pandemic. Cell 185, 447–456.

Hoffmann, M., Kleine-Weber, H., Schroeder, S., Krüger, N., Herrler, T., Erichsen, S., Schiergens, T.S., Herrler, G., Wu, N.H., Nitsche, A., Müller, M.A., Drosten, C., Pöhlmann, S., 2020a. SARS-CoV-2 cell entry depends on ACE2 and TMPRSS2 and is blocked by a clinically proven protease inhibitor. Cell 181, 271–280.

Hoffmann, M., Schroeder, S., Kleine-Weber, H., Müller, M.A., Drosten, C., Pöhlmann, S., 2020b. Nafamostat mesylate blocks activation of SARS-CoV-2: New treatment option for COVID-19. Antimicrob Agents Chemother 64, e00754–00720.

Jackson, C.B., Farzan, M., Chen, B., Choe, H., 2022. Mechanisms of SARS-CoV-2 entry into cells. Nat Rev Mol Cell Biol 23, 3–20.

Jiang, Y., van der Welle, J.E., Rubingh, O., van Eikenhorst, G., Bakker, W.A.M., Thomassen, Y.E., 2019. Kinetic model for adherent Vero cell growth and poliovirus production in batch bioreactors. Process Biochem 81, 156–164.

Ke, R., Zitzmann, C., Ho, D.D., Ribeiro, R.M., Perelson, A.S., 2021. In vivo kinetics of SARS-CoV-2 infection and its relationship with a person’s infectiousness. Proc Natl Acad Sci U S A 118, e2111477118.

Kissler, S.M., Fauver, J.R., Mack, C., Tai, C.G., Breban, M.I., Watkins, A.E., Samant, R.M., Anderson, D.J., Metti, J., Khullar, G., Baits, R., MacKay, M., Salgado, D., Baker, T., Dudley, J.T., Mason, C.E., Ho, D.D., Grubaugh, N.D., Grad, Y.H., 2021. Viral dynamics of SARS-CoV-2 variants in vaccinated and unvaccinated persons. N Engl J Med 385, 2489–2491.

Koch, J., Uckeley, Z.M., Doldan, P., Stanifer, M., Boulant, S., Lozach, P.Y., 2021. TMPRSS2 expression dictates the entry route used by SARS-CoV-2 to infect host cells. EMBO J 40, e107821.

Koizumi, Y., Ohashi, H., Nakajima, S., Tanaka, Y., Wakita, T., Perelson, A.S., Iwami, S., Watashi, K., 2017. Quantifying antiviral activity optimizes drug combinations against hepatitis C virus infection. Proc Natl Acad Sci U S A 114, 1922–1927.

Kreutzberger, A.J.B., Sanyal, A., Ojha, R., Pyle, J.D., Vapalahti, O., Balistreri, G., Kirchhausen, T., 2021. Synergistic block of SARS-CoV-2 infection by combined drug inhibition of the host entry factors PIKfyve kinase and TMPRSS2 protease. J Virol 95, e0097521.

Liu, J., Li, Y., Liu, Q., Yao, Q., Wang, X., Zhang, H., Chen, R., Ren, L., Min, J., Deng, F., Yan, B., Liu, L., Hu, Z., Wang, M., Zhou, Y., 2021. SARS-CoV-2 cell tropism and multiorgan infection. Cell Discov 7, 17.

Magnus, C., Rusert, P., Bonhoeffer, S., Trkola, A., Regoes, R.R., 2009. Estimating the stoichiometry of human immunodeficiency virus entry. J Virol 83, 1523–1531.

Meng, B., Ferreira, I.A.T.M., Abdullahi, A., Saito, A., Kimura, I., al., e., 2021. Altered TMPRSS2 usage by SARS-CoV-2 Omicron impacts infectivity and fusogenicity. Nature 603, 706–714.

Mulampaka, S.N., Dixit, N.M., 2011. Estimating the threshold surface density of Gp120-CCR5 complexes necessary for HIV-1 envelope-mediated cell-cell fusion. PLoS One 6, e19941.

Néant, N., Lingas, G., Le Hingrat, Q., Ghosn, J., Engelmann, I., Lepiller, Q., Gaymard, A., Ferré, V., Hartard, C., Plantier, J.C., Thibault, V., Marlet, J., Montes, B., Bouiller, K., Lescure, F.X., Timsit, J.F., Faure, E., Poissy, J., Chidiac, C., Raffi, F., Kimmoun, A., Etienne, M., Richard, J.C., Tattevin, P., Garot, D., Le Moing, V., Bachelet, D., Tardivon, C., Duval, X., Yazdanpanah, Y., Mentré, F., Laouénan, C., Visseaux, B., Guedj, J., 2021. Modeling SARS-CoV-2 viral kinetics and association with mortality in hospitalized patients from the French COVID cohort. Proc Natl Acad Sci U S A 118, e2017962118.

Ou, T., Mou, H., Zhang, L., Ojha, A., Choe, H., Farzan, M., 2021. Hydroxychloroquine-mediated inhibition of SARS-CoV-2 entry is attenuated by TMPRSS2. PLoS Pathog 17, e1009212.

Padmanabhan, P., Desikan, R., Dixit, N.M., 2020. Targeting TMPRSS2 and Cathepsin B/L together may be synergistic against SARS-CoV-2 infection. PLoS Comput Biol 16, e1008461.

Padmanabhan, P., Desikan, R., Dixit, N.M., 2022. Modeling how antibody responses may determine the efficacy of COVID-19 vaccines. Nat Comput Sci 2, 123–131.

Padmanabhan, P., Dixit, N.M., 2011. Mathematical model of viral kinetics in vitro estimates the number of E2-CD81 complexes necessary for hepatitis C virus entry. PLoS Comput Biol 7, e1002307.

Padmanabhan, P., Dixit, N.M., 2012. Viral kinetics suggests a reconciliation of the disparate observations of the modulation of claudin-1 expression on cells exposed to hepatitis C virus. PLoS One 7, e36107.

Padmanabhan, P., Dixit, N.M., 2015. Modeling suggests a mechanism of synergy between hepatitis C virus entry inhibitors and drugs of other classes. CPT Pharmacometrics Syst Pharmacol 4, 445–453.

Padmanabhan, P., Dixit, N.M., 2017. Inhibitors of hepatitis C virus entry may be potent ingredients of optimal drug combinations. Proc Natl Acad Sci U S A 114, E4524–E4526.

Padmanabhan, P., Garaigorta, U., Dixit, N.M., 2014. Emergent properties of the interferon-signalling network may underlie the success of hepatitis C treatment. Nature Commun 5, 3872.

Peacock, T.P., Brown, J.C., Zhou, J., Thakur, N., Sukhova, K., Newman, J., Kugathasan, R., Yan, A.W., Furnon, W., De Lorenzo, G., 2022. The altered entry pathway and antigenic distance of the SARS-CoV-2 Omicron variant map to separate domains of spike protein. bioRxiv.

Perelson, A.S., Ke, R., 2020. Mechanistic modeling of SARS-CoV-2 and other infectious diseases and the effects of therapeutics. Clin Pharmacol Ther, 10.1002/cpt.2160.

Ray, D., Le, L., Andricioaei, I., 2021. Distant residues modulate conformational opening in SARS-CoV-2 spike protein. Proc Natl Acad Sci U S A 118, e2100943118.

Sen, P., Saha, A., Dixit, N.M., 2019. You cannot have your synergy and efficacy too. Trends Pharmacol Sci 40, 811–817.

Ursache, R.V., Thomassen, Y.E., van Eikenhorst, G., Verheijen, P.J., Bakker, W.A., 2015. Mathematical model of adherent Vero cell growth and poliovirus production in animal component free medium. Bioprocess Biosyst Eng 38, 543–555.

Venugopal, V., Padmanabhan, P., Raja, R., Dixit, N.M., 2018. Modelling how responsiveness to interferon improves interferon-free treatment of hepatitis C virus infection. PLoS Comput Biol 14, e1006335.

Willett, B.J., Grove, J., MacLean, O.A., Wilkie, C., De Lorenzo, G., Furnon, W., Cantoni, D., Scott, S., Logan, N., Ashraf, S., Manali, M., Szemiel, A., Cowton, V., Vink, E., Harvey, W.T., Davis, C., Asamaphan, P., Smollett, K., Tong, L., Orton, R., Hughes, J., Holland, P., Silva, V., Pascall, D.J., Puxty, K., da Silva Filipe, A., Yebra, G., Shaaban, S., Holden, M.T.G., Pinto, R.M., Gunson, R., Templeton, K., Murcia, P.R., Patel, A.H., Klenerman, P., Dunachie, S., Haughney, J., Robertson, D.L., Palmarini, M., Ray, S., Thomson, E.C., 2022. SARS-CoV-2 Omicron is an immune escape variant with an altered cell entry pathway. Nat Microbiol 7, 1161–1179.

Xia, S., Wang, L., Zhu, Y., Lu, L., Jiang, S., 2022. Origin, virological features, immune evasion and intervention of SARS-CoV-2 Omicron sublineages. Signal Transduct Target Ther 7, 241.

Zhao, H., Lu, L., Peng, Z., Chen, L.L., Meng, X., Zhang, C., Ip, J.D., Chan, W.M., Chu, A.W., Chan, K.H., Jin, D.Y., Chen, H., Yuen, K.Y., To, K.K., 2021. SARS-CoV-2 Omicron variant shows less efficient replication and fusion activity when compared with delta variant in TMPRSS2-expressed cells. Emerg Microbes Infect 11, 277–283.

Zhuravel, S.V., Khmelnitskiy, O.K., Burlaka, O.O., Gritsan, A.I., Goloshchekin, B.M., Kim, S., Hong, K.Y., 2021. Nafamostat in hospitalized patients with moderate to severe COVID-19 pneumonia: a randomised Phase II clinical trial. EClinicalMedicine 41, 101169.

Zimmerman, M.I., Porter, J.R., Ward, M.D., Singh, S., Vithani, N., Meller, A., Mallimadugula, U.L., Kuhn, C.E., Borowsky, J.H., Wiewiora, R.P., Hurley, M.F.D., Harbison, A.M., Fogarty, C.A., Coffland, J.E., Fadda, E., Voelz, V.A., Chodera, J.D., Bowman, G.R., 2021. SARS-CoV-2 simulations go exascale to predict dramatic spike opening and cryptic pockets across the proteome. Nat Chem 13, 651–659.

